# *In Silico* method for identification of MHC class I-like molecules in whole genomes

**DOI:** 10.1101/046607

**Authors:** Peter Reinink, Ildiko van Rhijn

## Abstract

Mining of genomic data is a valuable tool for the discovery of orthologous genes of close or distant relatives of humans and mice. Here we describe a standardized search method for the MHC class I-like molecules CD1 and MR1 and apply it to 18 mammalian genomes.

## INTRODUCTION

Besides the well-known antigen presenting MHC class I and II molecules, also called the “classical MHC molecules” there are other antigen presenting molecules that are called MHC class I-like molecules based on close structural resemblance to MHC class I molecules. The most well-known MHC class I-like molecules are MR1 and CD1. In human this CD1 family exist of 5 isoforms; CD1a, CD1b, CD1c, CD1d, and CD1e^1^. Structurally, they are similar to the MHC class I molecules in that their antigen binding groove is composed of 2 domains, which are called α1 and an α2 domains. The third domain is an α3 domain which is linked to a transmembrane domain and a cytoplasmic tail. The major functional differences between MHC molecules and CD1 molecules are the ligands they bind. Whereas classical MHC molecules bind and present peptides to T cells, MR1 molecules bind small metabolites, and CD1 molecules bind lipids.

Among species there is a large discrepancy in the size of the CD1 locus and the number of genes for each isoform. For instance the mouse locus is smaller than the human locus and has only 2 genes. Both genes encode CD1d proteins and no genes for other isoforms are present in the mouse genome^2^. The canine locus on the other hand is much larger than the human locus. It has 4 functional CD1a genes and 5 CD1a pseudogenes, and furthermore there is 1 gene of each of the other isoforms^3–5^.

This difference in size and composition of the CD1 families among species makes it difficult to identify and assign these MHC class I-like molecules. We developed and validated a method that is particularly well suited for MHC class I-like molecules.

## METHODS

### Genomic databases

For every species examined in this study the most recent Soft-masked assembly sequence was downloaded from the UCSC genome browser (http://hgdownload.soe.ucsc.edu/downloads.html), (Table 1)^6^. These assembly files were reformatted to match the requirement of the makeblastdb function from BLAST+^7^ for generating nucleotide databases (i.e. a single line of sequence after the header).

**Table 1:**
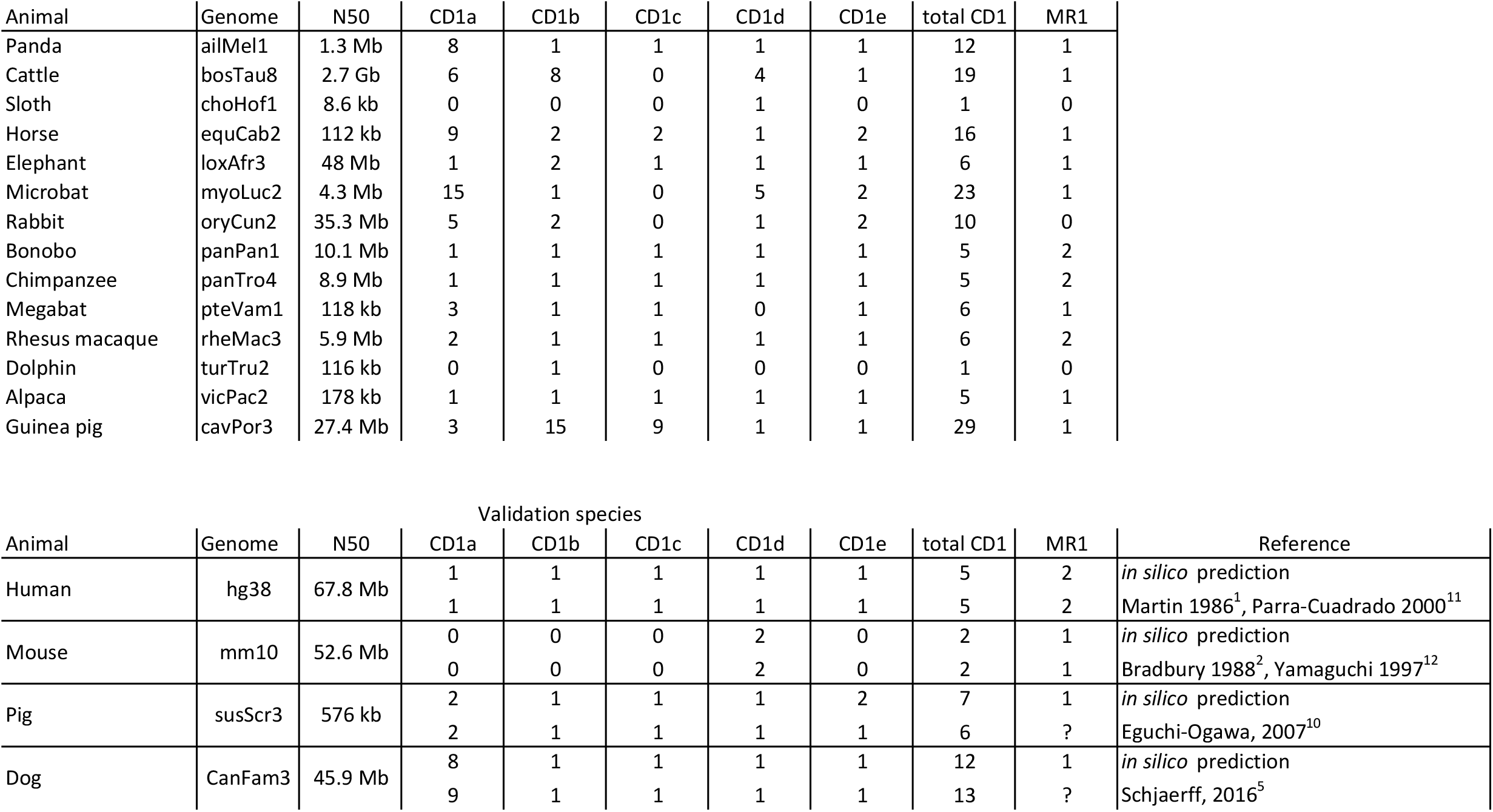
Overview of CD1 and MR1 orthologs in mammals. Overview of the number of CD1 and MR1 genes found in the species tested in this study. The N50 size is the length such that 50% of the assembled genome lies in blocks of the N50 size or longer. For human, mouse, dog, and pig, which we used as validation species, the previously published numbers of genes are also shown.

### Alignment-based searches

To find homologs of non-classical MHC molecules we used a Basic Local Alignment Search Tool for nucleotide queries (BLASTN) with default parameters. For post-BLASTN analysis the output format was set to output a tabular output. As input a fasta file with the nucleotide sequences of the second exons and the third exons of human non-classical MHC molecules was used, which code for the α1 and α2 domains, respectively.

### Post-BLASTN analysis

BLAST hits were filtered on their alignment length (>60 bp), e-value (<0.1) and on being located on a chromosome or a genomic scaffold of >4500 bp. These parameters were determined experimentally and subsequently used for all analyses we report here. If multiple BLAST hits overlapped, only the first BLAST hit was used for further analysis. We call the results of this initial filtering step “unique BLAST hits”. Furthermore, BLAST hits that were shared among the results obtained with exon2 and exon3 were excluded. BLAST hits were ordered according to chromosomal location. A combination of an exon2 and an exon3 BLAST hit was called a “BLAST pair” (i.e. a α1-α2 domain combination) if the distance between them was smaller than 2400 base pairs and they were located on the same strand.

### Alignment and clustering

Exon2-exon3 BLAST hits were merged and the merged sequences were aligned with a collection of cDNA sequences of the exon 2 and exon 3 sequences of the known human non-classical MHC molecules. This collection contained murine H2-M3 and human CD1a, CD1b, CD1c, CD1d, CD1e, MR1, MICA, MICB, and classical MHC class I molecules. The alignment was performed with MUSCLE^8^. These alignments were used as input for ClustalW Phylogeny^9^ to create a rooted dendrogram which was subsequently used to assign the correct name to BLAST pairs via clustering with an Unweighted Pair Group Method with Arithmetic Mean (UPGEMA) algorithm.

## RESULTS

### Method validation

Before the algorithm was applied to mammalian species in which the CD1 molecules had never been studied before, we first determined the sensitivity and accuracy of the developed method. To do this, the second and third exons of human CD1a, CD1b, CD1c and CD1d were used as input for the BLASTN search in the human genome. The BLASTN result showed a total of 189 non unique BLAST hits. These BLAST hits were filtered as described in the methods section. After this filtering step the number was reduced to 10 unique BLAST hits that were all located on chromosome 1, were the human CD1 locus is known to be located. Of the ten BLAST hits on chromosome 1, five BLAST pairs could be formed. The resulting five BLAST pairs represented not only the four CD1 molecules that were used as input for the initial search but included CD1e as well, which is the fifth member of the human CD1 family (Figure 1a, Table 2).

**Figure 1:**
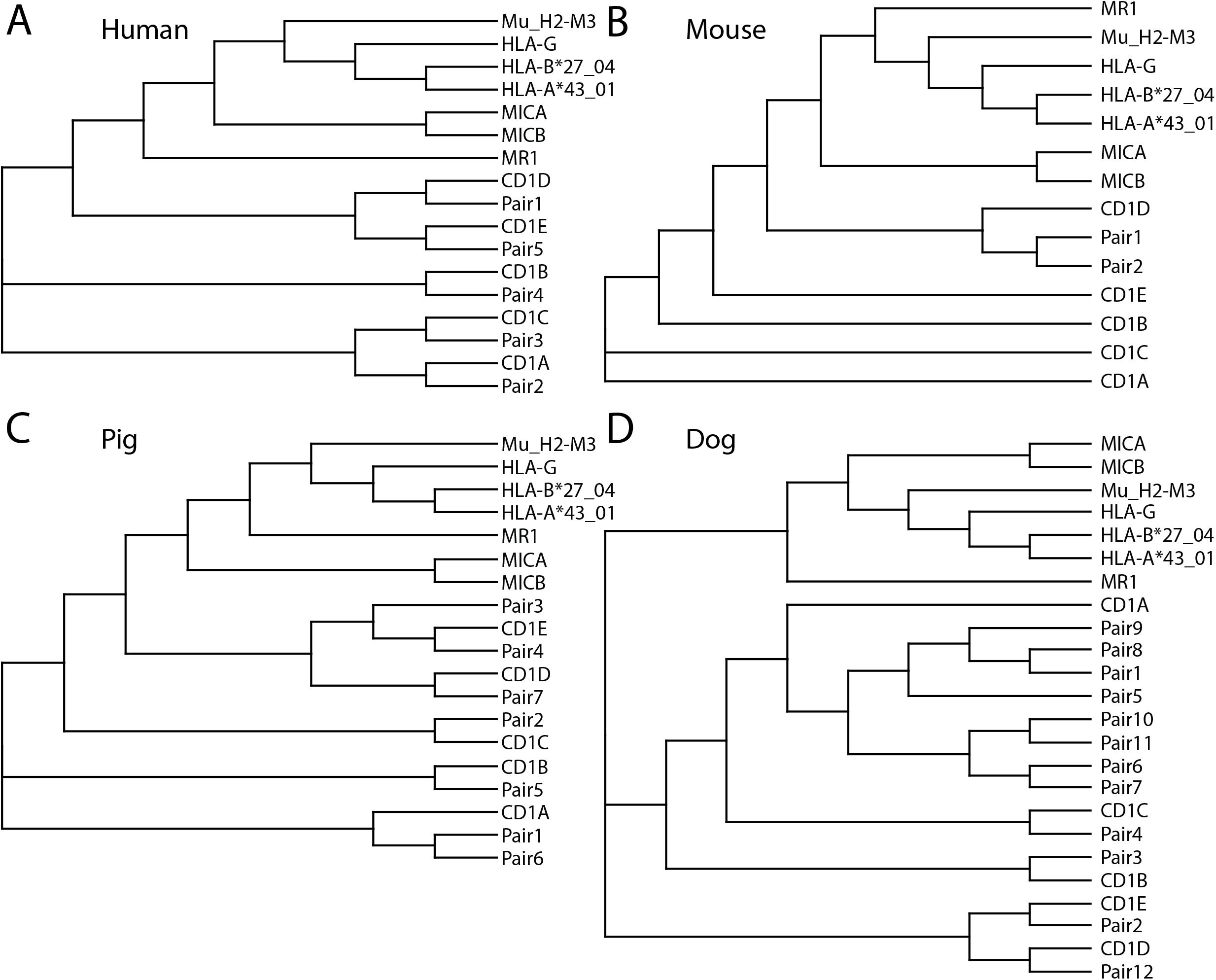
Method validaton. BLAST pairs that resulted from searches in the A) human, B) mouse, C) porcine and D) canine genome were aligned with the combined α1 and α2 domains of the human CD1 family members, MR1, MICA, MICB, HLA-G, HLA-A, HLA-B and Murine M2-H3. The alignment is shown as a rooted dendrogram.

**Table 2:** Locations of BLAST pairs and locations of CD1 molecules that form the basis of Figure 1. BLAST pairs that resulted from searches, including the locations of the individual BLAST hits, are shown. For human and mouse the annotated locations of the known CD1 were added.

| Species (genome) | Pair | Chr | start | stop | Hit 1 |  | Hit 2 |  | Strand | CD1 genes annotated in ENSEMBL |  |  |  |  |
| --- | --- | --- | --- | --- | --- | --- | --- | --- | --- | --- | --- | --- | --- | --- |
|  |  |  |  |  | start | stop | start | stop |  | Name | Chr | start | stop | strand |
| Human (HG38) | P1 | chr1 | 158181485 | 158182137 | 158181485 | 158181608 | 158182032 | 158182137 |  | CD1D | chr1 | 158179947 | 158184896 |  |
|  | P2 | chr1 | 158255084 | 158256282 | 158255084 | 158255350 | 158256004 | 158256282 |  | CD1A | chr1 | 158254137 | 158258269 |  |
|  | P3 | chr1 | 158291149 | 158292346 | 158291149 | 158291313 | 158292084 | 158292346 |  | CD1C | chr1 | 158289786 | 158293630 |  |
|  | P4 | chr1 | 158329852 | 158331059 | 158329852 | 158330128 | 158330884 | 158331059 | reverse | CD1B | chr1 | 158327951 | 158331531 | reverse |
|  | P5 | chr1 | 158354448 | 158355522 | 158354448 | 158354585 | 158355349 | 158355522 |  | CD1E | chr1 | 158353696 | 158357553 |  |
| Mouse (mm10) | P1 | chr3 | 86986988 | 86987821 | 86986988 | 86987241 | 86987543 | 86987821 |  | Cd1D2 | chr3 | 86986551 | 86989780 |  |
|  | P2 | chr3 | 86998072 | 86998904 | 86998072 | 86998350 | 86998652 | 86998904 | reverse | Cd1D1 | chr3 | 86995834 | 86999441 | reverse |
| Pig (susScr3) | P1 | chr4 | 99906355 | 99907575 | 99906355 | 99906559 | 99907297 | 99907575 |  |  |  |  |  |  |
|  | P2 | chr4 | 99930347 | 99932095 | 99930347 | 99930520 | 99931817 | 99932095 |  |  |  |  |  |  |
|  | P3 | chr4 | 99969612 | 99969961 | 99969612 | 99969718 | 99969885 | 99969961 | reverse |  |  |  |  |  |
|  | P4 | chr4 | 100359279 | 100361016 | 100359279 | 100359401 | 100360908 | 100361016 | reverse |  |  |  |  |  |
|  | P5 | chr4 | 100380755 | 100383144 | 100380755 | 100381031 | 100383023 | 100383144 | reverse |  |  |  |  |  |
|  | P6 | chr4 | 100410977 | 100412203 | 100410977 | 100411240 | 100412005 | 100412203 | reverse |  |  |  |  |  |
|  | P7 | chr4 | 100440602 | 100441438 | 100440602 | 100440878 | 100441191 | 100441438 | reverse |  |  |  |  |  |
| Dog (canFam3) | p1 | chrUn_JH374180 | 2251 | 3472 | 2251 | 2423 | 3194 | 3472 |  |  |  |  |  |  |
|  | p2 | chr38 | 23296481 | 23297474 | 23296481 | 23296611 | 23297375 | 23297474 | reverse |  |  |  |  |  |
|  | p3 | chr38 | 23318380 | 23319500 | 23318380 | 23318648 | 23319235 | 23319500 | reverse |  |  |  |  |  |
|  | p4 | chr38 | 23329381 | 23330954 | 23329381 | 23329518 | 23330705 | 23330954 | reverse |  |  |  |  |  |
|  | p5 | chr38 | 23348267 | 23349484 | 23348267 | 23348540 | 23349312 | 23349484 | reverse |  |  |  |  |  |
|  | p6 | chr38 | 23363867 | 23365085 | 23363867 | 23364145 | 23364871 | 23365085 | reverse |  |  |  |  |  |
|  | p7 | chr38 | 23383469 | 23384673 | 23383469 | 23383747 | 23384485 | 23384673 | reverse |  |  |  |  |  |
|  | p8 | chr38 | 23404220 | 23405452 | 23404220 | 23404497 | 23405276 | 23405452 | reverse |  |  |  |  |  |
|  | p9 | chr38 | 23424873 | 23426091 | 23424873 | 23425145 | 23425888 | 23426091 | reverse |  |  |  |  |  |
|  | p10 | chr38 | 23437848 | 23439066 | 23437848 | 23438129 | 23438891 | 23439066 | reverse |  |  |  |  |  |
|  | p11 | chr38 | 23455806 | 23457004 | 23455806 | 23456084 | 23456925 | 23457004 | reverse |  |  |  |  |  |
|  | p12 | chr38 | 23491100 | 23491712 | 23491100 | 23491376 | 23491485 | 23491712 | reverse |  |  |  |  |  |

After this initial validation, we applied our method to other animals of which the CD1 locus is well studied. We started with mouse (mm10), which is known to have 2 CD1d homologs and no CD1a, CD1b, CD1c and CD1e homologs^2^. The result of a search with the human CD1a, CD1b, CD1c, and CD1d exon2 and exon3 gave 141 non unique BLAST hits that after filtering were reduced to four. These four BLAST hits could be combined to 2 pairs. Both grouped with HuCD1d (Figure 1b) and their chromosomal location and orientation overlap with those of the murine *Cd1d1* and *Cd1d2* genes (Table 2).

The third test was on pig (susScr3), which is known to have 6 CD1 genes: two CD1a homologs and one homolog for each of the other CD1 genes^10^. 141 Non unique BLASTN hits were reduced to 14 unique BLAST hits after filtering and formed 7 BLAST pairs. By aligning the 7 BLAST pairs to the Human CD1 sequences a possible new CD1e homolog was found that was not described before (Figure 1c). This possible new homolog is located in the CD1 locus of the pig (Table 2).

As a final validation step, our search strategy was applied to the canine genome assembly (canFam3). The canine CD1 locus is located on chromosome 38. Currently, 13 canine CD1 genes are known: 9 CD1a homologs, and 1 homolog for each of the other CD1 isoforms^3,5^. Of the 163 non unique BLAST hits, 25 BLAST hits past the filtering step. These resulted in 12 BLAST pairs, of which 8 BLAST pairs clustered with CD1a and one BLAST pair for each the other human CD1 genes (Figure 1d). One of the 8 BLAST pairs that cluster with CD1a is located on the unassigned contig chrUn_JH374180 and all other BLAST pairs are located on chromosome 38 (Table 2). Please note a discrepancy between the numbers of BLAST pairs found via our method (12) and the number of canine CD1 genes that has been reported by Schjaerff et al. (13)^5^.

Although most known CD1 homologs were found during the validation of our method in the human, mouse, pig, and dog genome, we would like to point out that our method does not discriminate between functional and non-functional genes. BLAST pairs identified with our method can be pseudogenes as well as functional genes.

### CD1 genes in other mammalian species

Next, we proceeded to apply our validated method to the following mammalian genomes: cow (bosTau8), horse (equCab2), African elephant (loxAfr3), bonobo (panPan1), chimpanzee (panTro4), alpaca (vicPac2), rhesus macaque (rheMac3), dolphin (turTru2), sloth (choHof1), panda (ailMel1), megabat (pteVam1), microbat (myoLuc2), guinea pig (canPor3), and rabbit (oryCun2). For all of these species 1 or more CD1 homologs were found (Figure 2, Table 1, Table S1). Our data show that in the tested mammals, multiplication of CD1a is very common, and multiplication of CD1b, CD1c, and CD1d is more common than multiplications of CD1e. To confirm the unexpectedly high number of CD1 homologs in guinea pig an additional BLASTN search was performed with the human CD1a α3 domain (exon 4). For all 29 BLAST pairs an α3 domain was found within 600 base pairs downstream of the α2 domain. This confirms that the α1-α2 BLAST pairs that were initially identified are likely to be part of CD1 paralogs. However, even having a correct combination of α1, α2, and α3 domain provides no information concerning the functionality of the CD1 gene. One way to assess potential functionality of a gene is to determine whether it can give rise to an open reading frame. To do this, we compared the 29 BLAST pairs that we found with predicted ORFs in the same chromosomal region in ENSEMBL (<u>www.ensembl.org</u>). Among the 29 BLAST pairs we found, 15 were part of predicted ORFs in ENSEMBL. Of those 15 predicted ORFs, 10 were annotated as CD1 gene including isoform, and 5 were annotated as CD1, but the isoform was not determined. Based on grouping with the human isoforms, we were able to assign these five CD1 genes of undetermined isoform as 3 CD1b homologs and 2 CD1c homologs. The 14 BLAST pairs that were not part of predicted ORFs can either be pseudogenes or functional genes for which the ORF-prediction algorithm could not predict an ORF. For comparison between our method applied to the guinea pig genome and annotated ORFs in ENSEMBL, a tree combining the ORFs and the pairs resulting from our method was generated (Figure 3). This result shows the power of this new method of homology searching.

**Figure 2:**
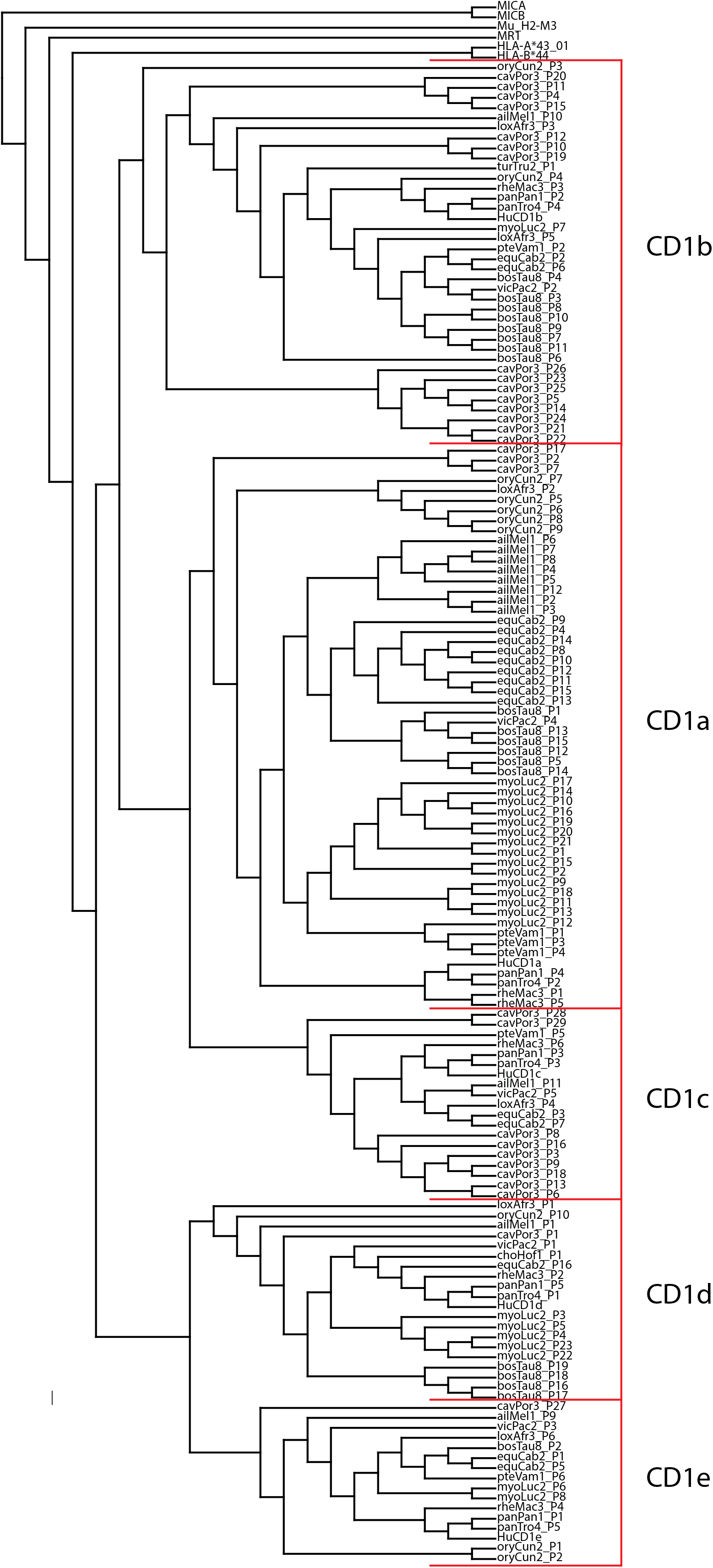
CD1 genes in other mammalian species. BLAST pairs that resulted from CD1-targeted searches in the indicated species were aligned with the combined α1 and α2 domains of the human CD1 family members, MR1, MICA, MICB, HLA-A, HLA-B and Murine M2-H3. The alignment is shown as a rooted dendrogram.

**Figure 3:**
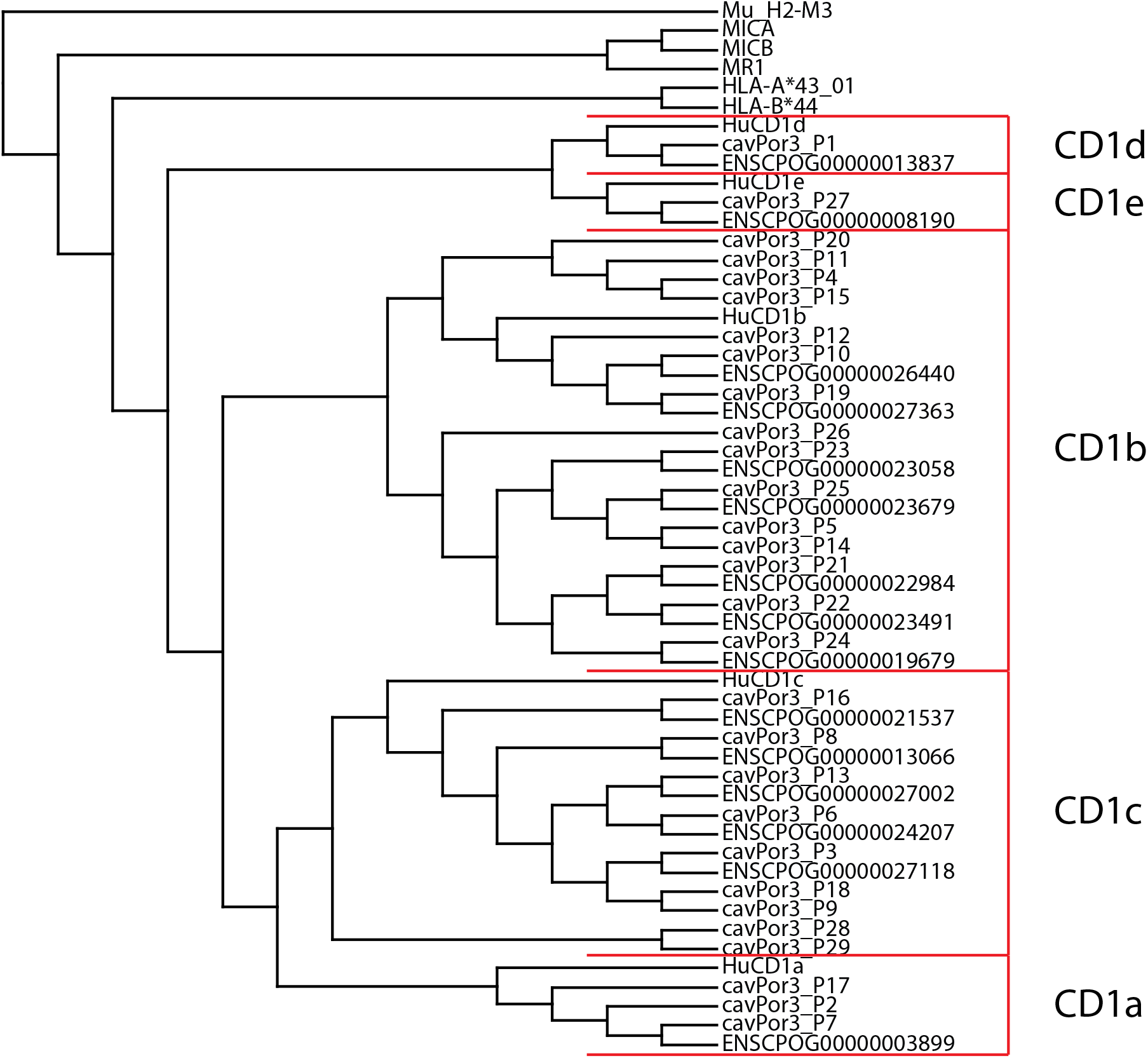
Comparison between BLAST search and known ORFs in the guinea pig CD1 locus. Rooted dendrogram of the BLAST pairs resultng from our CD1-targetng search in the guinea pig genome and known ORFs in the guinea pig genome as annotated in ENSEMBL.

### MR1 genes in mammalian species

MR1 genes are not known to exist as larger groups of genes or in varieties analogous to the CD1 isoforms. To study the presence of MR1 in various species we applied our method to MR1. We performed BLASTN searches with exon3 and exon4 of human MR1 in the same mammalian genomes as we used for the CD1 searches. For sloth, rabbit, and dolphin no MR1 BLAST hits were found. However, we noted that the sloth and dolphin genomes are among the genomes with the smallest N50 value, which is an indicator of how big the chunks of sequence are that make up the genome assembly. It is possible that the low N50 value in sloth and dolphin led to potential false negative results using our BLAST-based method. The rabbit genome assembly has a higher N50 value, but no MR1 gene was found. We consider it possible, but not proven, that rabbits do not have a gene for MR1. For most other mammals 1 MR1 BLAST pair was found except for the primates, there 2 MR1 BLAST pairs were found (Table 1). For humans 1 BLAST pair overlaps with the MR1 gene the other BLAST pair overlaps with the known pseudo gen RP11-46A10.6 (ENSG00000251520). This gene is annotated as a “pseudogene similar to part of major histocompatibility complex, class I-related MR1”. All these BLAST pairs cluster with the human MR1 gene, forming an interspecies groups distinct from CD1 (Figure 4, Table S2). Our MR1 results show that this *in silico* prediction method can also be used for MHC class I-like molecules other than CD1.

**Figure 4:**
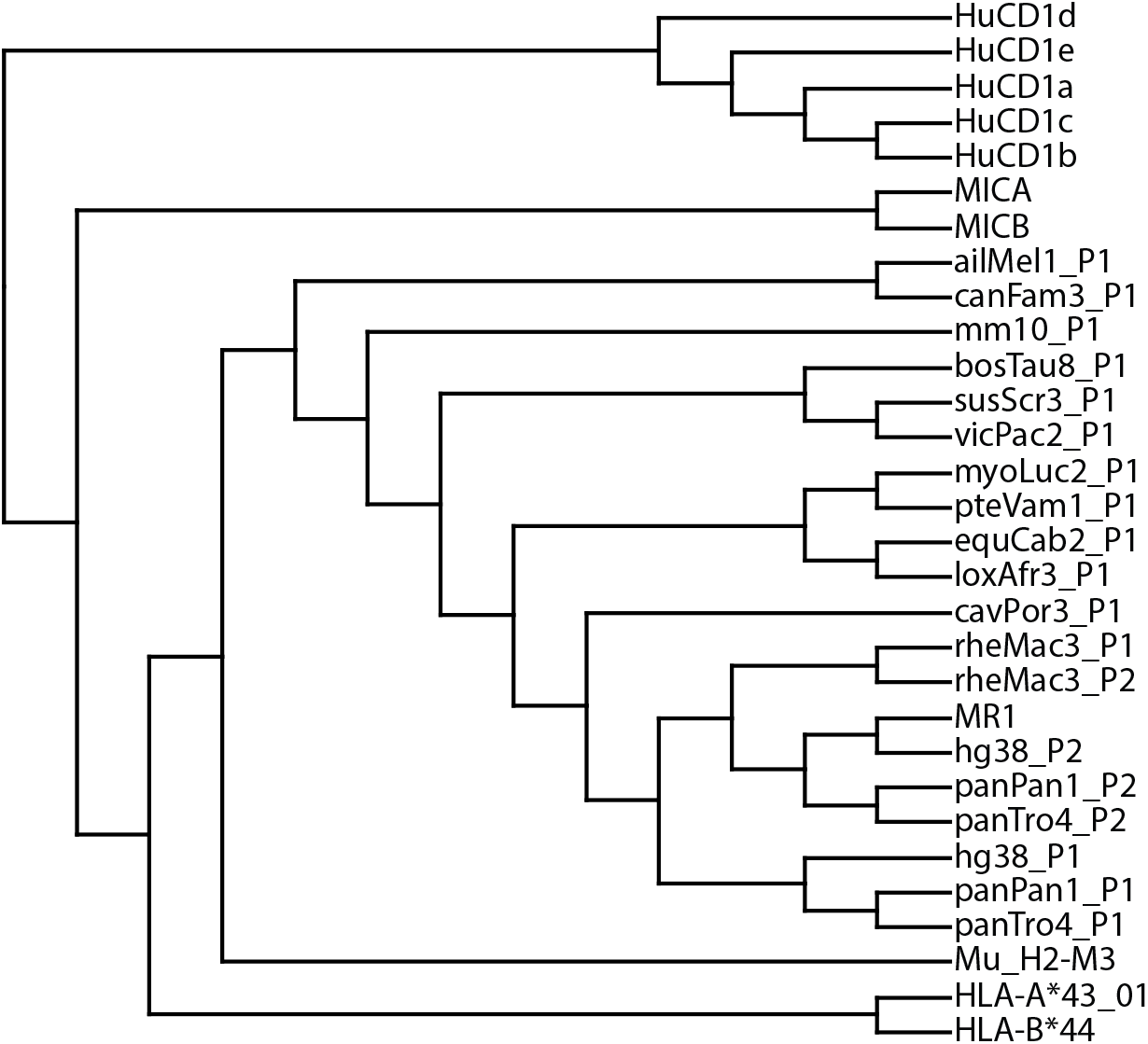
MR1 genes in mammals. BLAST pairs resultng from an MR1-targeted search for all indicated species were aligned with the combined α1 and α2 domains of the human CD1 family members, MR1, MICA, MICB, HLA-A, HLA-B and Murine M2-H3. The alignment is shown as a rooted dendrogram.

## DISCUSSION

This study is the first attempt to describe MHC class I like molecules of many mammals, including ones that have not previously been studied, in an *in silico* manner. The automated BLAST method we describe here consists of a blast search with the sequences of exon2 and exon3 which together form the antigen binding groove. This step is followed by a filtering step and pairing of BLAST hits that originate from different exons. The results of our method applied to genomes with well-studied CD1 loci (human, mouse, pig, dog) are highly consistent with the published data. Overall, when applied to species that have not been studied before, we find a highly variable number of CD1 genes and only one or two MR1 genes. The variability in size of the CD1 loci is in line with the sizes of the CD1 family indicated by other methods like cDNA cloning.

This method performs best with well assembled genomes like the human or mouse genome. The more incomplete the assembly of the genome is, the more difficult it is to find all homologs. In incompletely assembled genomes the following problems can arise: genes can be missed because the chromosome size is too small to pass the filter step or both exons are not located on the same DNA fragment. Furthermore, misassemblies of repeated sequences, collapses of repeated regions, and unmerged overlaps due to polymorphisms resulting in artificial duplications can occur in incompletely assembled and curated genomes. Since this method does not include splicing information and only searches with the α1 and α2 domains, no prediction can be made on the functionality of the resulting pairs. The BLAST pairs resulting from the BLAST search can be either functional genes or pseudogenes.

The question has been raised whether the distinction of the five CD1 isoforms CD1a, CD1b, CD1c, CD1d, and CD1e is merely based on the human situation with its five different CD1 genes, or on real biological differences that justify exactly five isoforms. The results we describe here support the idea that there are exactly five groups of CD1 genes. We were open to the possibility of identifying mammalian CD1 genes that do not cluster with the known isoforms, and in fact, our results included CD1e isoforms, despite the fact that the search was performed with CD1a, CD1b, CD1c, and CD1d only. However, among the current set of mammalian genomes we studied we found no evidence for additional CD1 isoforms or single CD1 genes that do not cluster with one of the existing five isoforms.^11,12^

